# Chromatin Functional States Correlate with HIV Latency Reversal in Infected Primary CD4^+^ T cells

**DOI:** 10.1101/242958

**Authors:** Emilie Battivelli, Matthew S. Dahabieh, Mohamed Abdel-Mohsen, J. Peter Svensson, Israel Tojal Da Silva, Lilian B. Cohn, Andrea Gramatica, Steven Deeks, Warner Greene, Satish K. Pillai, Eric Verdin

## Abstract

Human immunodeficiency virus (HIV) infection cannot be cured due to a small reservoir of latently infected CD4^+^ T cells in treated patients. The “shock and kill” approach proposes to eliminate the reservoir by inducing its activation and the direct or indirect killing of infected cells. Current latency reversing agents (LRAs) do not reduce the viral reservoir in treated patients.

We use a novel dual-fluorescent HIV reporter to identify and purify latent cells, and to determine the fraction of latent cells that undergo viral reactivation after infection of primary CD4^+^ T cells. Unexpectedly, LRAs reactivate less than 5% of latent proviruses. Analysis of HIV integration sites from induced and non-induced latent populations reveals distinct provirus integration sites between these two populations in terms of chromatin functional states.

These findings challenge “shock and kill”, and suggest the need of more potent LRAs in combination with immunomodulatory approaches to eradicate HIV reservoir.

## INTRODUCTION

Antiretroviral therapy (ART) has transformed HIV infection from a deadly disease into a chronic lifelong condition, saving millions of lives. However, ART interruption leads to rapid viral rebound within weeks due to the persistence of proviral latency in rare, long-lived resting CD4^+^ T cells and, to an unknown extent, in other cell populations. HIV latency is defined as the presence of a transcriptionally silent but replication-competent proviral genome that allows infected cells to evade both immune clearance mechanisms and ART.

A possible approach to purging latent HIV is the “shock and kill” strategy, which consists of forcing the reactivation of latent proviruses (“shock” phase) with the use of latency reversing agents (LRAs), while maintaining ART to prevent *de novo* infections. Subsequently, reactivation of HIV expression would expose such cells (shocked cells) to killing by viral cytopathic effects and immune clearance (“kill” phase).

A variety of LRAs have been explored *in vitro* and *ex vivo,* with only a few candidates being advanced to testing in pilot human clinical trials for their ability to reverse HIV latency. Direct proof-of-concept of histone deacetylase inhibitors (HDACi: vorinostat, panobinostat, romidepsin, and disulfiram) in clinical studies has shown increases in cell-associated HIV RNA production and/or plasma viremia after *in vivo* administration (Archin, Liberty, et al., 2012; Elliott et al., 2015; Elliott et al., 2014; Rasmussen et al., 2014; Sogaard et al., 2015). However, none of these interventions alone has succeeded in significantly reducing the size of the latent HIV reservoir (Rasmussen & Lewin, 2016).

Several obstacles can explain the failure of LRAs, as reviewed in (Margolis, Garcia, Hazuda, & Haynes, 2016; Rasmussen, Tolstrup, & Sogaard, 2016). However, the biggest challenge to date is our inability to accurately quantify the size of the reservoir. The absolute quantification (number of cells) of the latent reservoir *in vivo (and ex vivo)* has been technically impossible. The most sensitive, quickest, and easiest assays to measure the prevalence of HIV-infected cells are PCR-based assays that quantify total or integrated HIV DNA or RNA transcripts. However, these assays overestimate the number of latently infected cells due to the predominance of defective HIV DNA genomes *in vivo* (Bruner et al., 2016; Ho et al., 2013). The gold-standard assay to measure the latent reservoir is a viral outgrowth assay (VOA), which is neither quick nor easy, and consists of quantifying the number of resting CD4^+^ T cells that produce infectious virus after a single round of maximum *in vitro* T-cell activation. After several weeks of culture, viral outgrowth is assessed by an ELISA assay for HIV-1 p24 antigen or a PCR assay for HIV-1 RNA in the culture supernatant. However, the number of latently infected cells detected in the VOA is 300-fold lower than the number of resting CD4^+^ T cells that harbor proviruses detectable by PCR. This reliance on a single round of T-cell activation likely underestimates the viral reservoir for several reasons:

a. The stochastic nature of HIV activation (Dar et al., 2012; Ho et al., 2013; Singh, Razooky, Cox, Simpson, & Weinberger, 2010; Weinberger, Burnett, Toettcher, Arkin, & Schaffer, 2005). Two elegant studies show that the discovery of intact non-induced proviruses indicates that the size of the latent reservoir may be much greater than previously thought. The authors estimate that the number may be at least 60 fold higher than estimates based on VOA (Ho et al., 2013; Sanyal et al., 2017). One important point being highlighted with their work and with other’s (Chen, Martinez, Zorita, Meyerhans, & Filion, 2016) is the heterogeneous nature of HIV latency.
b. The ability of defective proviruses to be transcribed and translated *in vivo* (Pollack et al., 2017). This study shows that, although defective proviruses cannot produce infectious particles, they express viral RNA and proteins, which can be detectable by any p24 antigen or PCR assay used for the reservoir-size quantification.

Thus, current assays misestimate the absolute number of latently infected cells (true viral reservoir size) *in vivo and ex vivo,* and the size of HIV reservoir is still to be determined. Therefore, it has been difficult to judge the potential of LRAs in *in vitro* (latency primary models), *ex-vivo* (patients’ samples) and *in vivo* (clinical trial) experiments.

In addition, HIV latency is a complex, multi-factorial process (reviewed in (Dahabieh, Battivelli, & Verdin, 2015)). Its establishment and maintenance depend, in part, on: (a) viral factors, such as integrase that specifically interacts with cellular proteins, including LEDGF, (b) *trans-acting* factors (e.g., transcription factors) and their regulation by the activation state of T cells and the environmental cues that these cells receive, and (c) *cis*-acting mechanisms, such as the site of integration of the virus into the genome and the local chromatin environment.

Lack of knowledge about the viral reservoir has obstructed our ability to understand the relationship between viral integration and viral transcription. Several groups have studied this relationship and have reported conflicting data. While two studies failed to find a significant role of integration sites in regulating the fate of HIV infection (Dahabieh et al., 2014; Sherrill-Mix et al., 2013), other studies found that the HIV integration site does affect both the entry into latency (Chen et al., 2016; Jordan, Bisgrove, & Verdin, 2003; Jordan, Defechereux, & Verdin, 2001), and the viral response to LRAs (Chen et al., 2016). Thus, the correlation between integration sites and the fate of HIV-1 infection remains unclear.

In this study, we used a new dual color reporter virus, HIV_GKO_, to investigate the reactivation potential of various LRAs in pure latent populations. Although the quantification of cell-associated HIV RNA of HIV_GKO_ latently infected cells are consistent with results from patients’ samples, the various tested LRAs only reactivate virus within a small fraction (< 5%) of purified latently infected cells. To understand why some latent proviruses do not reactivate, we sequence HIV integration sites from induced and non-induced infected populations, and show that genomic localization and chromatin context of the integration site affects the fate of HIV infection and the reversal of viral latency.

## RESULTS

### A second-generation dual-fluorescence HIV-1 reporter (HIV_GKO_) to study latency

Our laboratory recently reported the development of a dual-labeled virus (DuoFluoI) in which eGFP is under the control of the HIV-1 promoter in the 5'LTR and mCherry is under the control of the cellular elongation factor 1 alpha promoter (EF1α) (Calvanese, Chavez, Laurent, Ding, & Verdin, 2013). However, we noted that the model was limited by a modest number of latently infected cells (<1%) generated regardless of viral input, as well as a high proportion of productively infected cells in which the constitutive promoter EF1α was not active (GFP+, mCherry-).

To address these issues, which we suspected were due to recombination between the 20–30-bp regions of homology at the N- and C-termini of the adjacent fluorescent proteins (eGFP and mCherry) (Salamango, Evans, Baluyot, Furlong, & Johnson, 2013), we generated a new version of dual-labeled virus (HIV_GKO_), containing a codon-switched eGFP (csGFP) and a distinct, unrelated fluorescent protein mKO2 under the control of EF1α (Figure 1A). First, titration of HIV_GKO_ input revealed that productively and latently infected cells increased proportionately as the input virus increased (Figure 1B), unlike the original DuoFluo I. Second, comparison of primary CD4^+^ T cells infected with HIV_GKO_ or the original DuoFluoI revealed an increase in double-positive (csGFP+ mKO2+), productively infected cells in HIV_GKO_ infected cells (Figure 1C). A small proportion of csGFP+ mKO2-cells were still visible in HIV_GKO_ infected cells. We generated a HIV_GKO_ virus lacking the U3 promoter region of the 3'LTR (DU3-GKO), resulting in an integrated virus devoid of the 5’ HIV U3 region. This was associated with a suppression of HIV transcription and an inversion of the latency ratio (ratios latent/productive = 0.34 for HIV_GKO-WT-LTR_ and 8.8 for HIV_GKO-∆U3-3’LTR_ - Figure 1D). Finally, to further characterize the constituent populations of infected cells, double-negative cells, latently and productively infected cells were sorted using FACS and analyzed for viral mRNA and protein content. (Figures 1E, F). As expected, productively infected cells (csGFP+ mKO2+) expressed higher amounts of viral mRNA and viral proteins, but latently infected cells (csGFP-mKO2+) had very small amounts of viral mRNA and no detectable viral proteins.

**Fig. 1.**
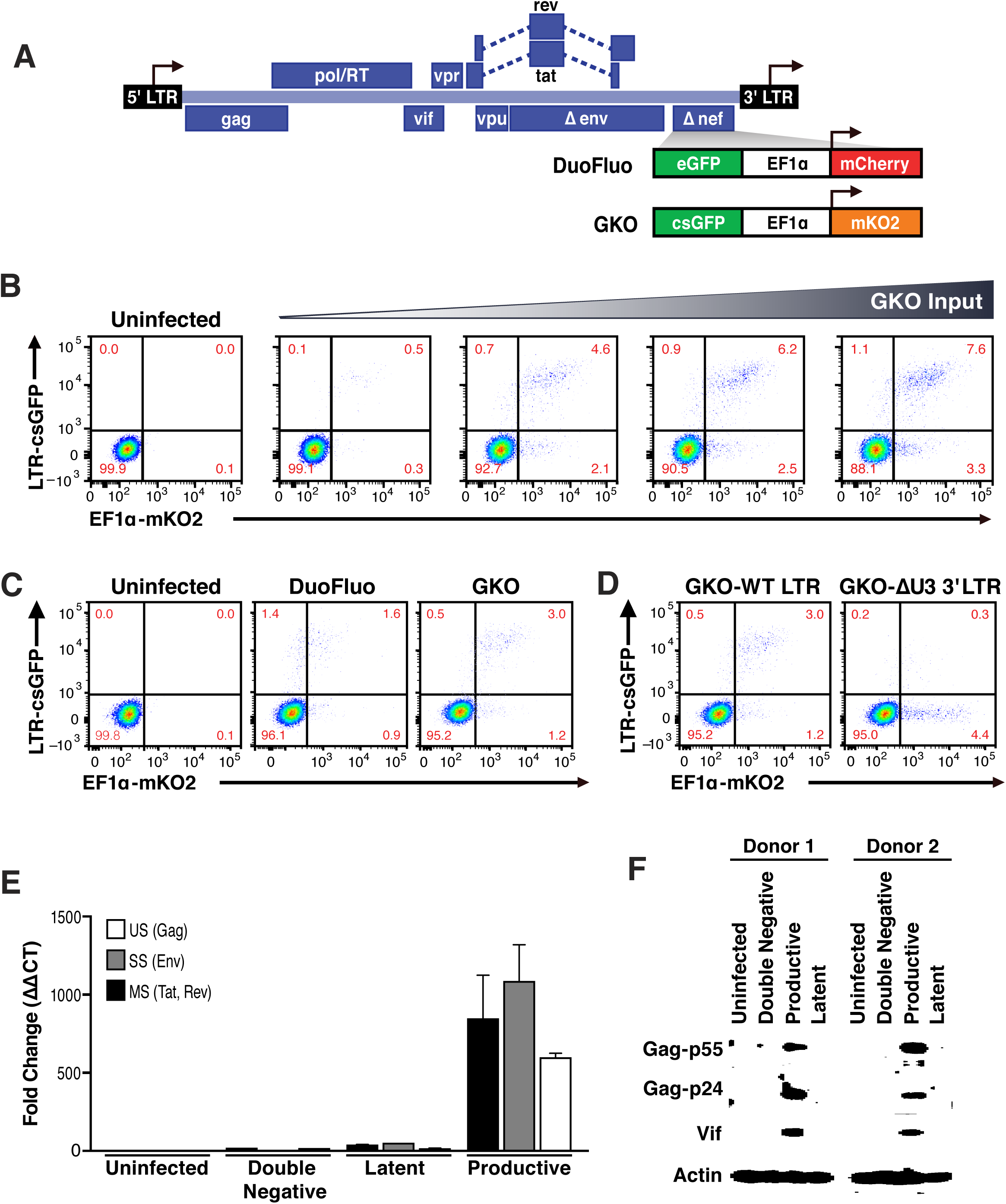
Second generation of dual-fluorescence HIV-1 reporter, HIV_GKO_ to quantify stable latency. (A) Schematic representation of first (top: HIV_DuoFluoI_) and second generation (bottom: HIV_GKO_) of dual-labeled HIV-1 reporters. (B, C) Primary CD4^+^ T cells were activated with αCD3/CD28 beads + 20 U/mL IL-2 for 3 days before infection. At 4 days post-infection, double-negative, productively infected, and latently infected cells were sorted out, and (B) the total RNA isolated from each population was subjected to Taqman RT-qPCR analysis. Unspliced (US), singly spliced (SS), and multiply spliced (MS) HIV-1 mRNAs were quantified relative to cellular GAPDH. (C) Western blot analysis of each population. (D) Comparison of HIV_DuoFluoI_ and HIV_GKO_ infection profiles by flow cytometry in activated primary CD4^+^ T-cells (4 days post-infection). Cells were treated as in (B). (E) Representative experiment of HIV_GKO_ virus titration in activated primary CD4^+^ T cells (4 days post-infection). Cells treated as in (B) were infected with different amounts of HIV_GKO_ (input, ng/p24) and analyzed by flow cytometry 4 days post-infection. (F) Comparison of GKO-WT-LTR and GKO-∆U3 3'LTR infection profiles by flow cytometry in cells treated as in (B).

Taken together, the second-generation of dual-fluorescence reporter, HIV_GKO_, is able to more accurately quantify latent infections in primary CD4^+^ T cells, and allows for the identification and purification of a much larger number of latently infected cells. By recording the read-out using flow cytometry, we can determine infection and HIV productivity of individual cells and simultaneously control for cell viability.

### LRA efficacy in patient samples is predicted by activity in HIV_GKO_ latently infected cells

Next, we evaluated the reversal of HIV_GKO_ latently infected primary CD4^+^ T cells by LRAs, and compared it with the reversal of latency in HIV infected individuals cells using the same LRAs. We tested the most well described mechanistically distinct LRAs to asses their latency reactivation potential: (a) the histone deacetylase inhibitor (HDACi) panobinostat (Rasmussen, Tolstrup, Winckelmann, Ostergaard, & Sogaard, 2013), (b) the bromodomain-containing protein 4 (BRD4) inhibitor JQ1, which acts through positive transcription elongation factor (P-TEFb) (Banerjee et al., 2012; Boehm et al., 2013; Filippakopoulos et al., 2010; Li, Guo, Wu, & Zhou, 2013; Zhu et al., 2012), and (c) a PKC activator, bryostatin-1 (del Real et al., 2004; Mehla et al., 2010). Viral reactivation mediated by these LRAs was compared to CD4^+^ T cells treated with αCD3/CD28 (Spina et al., 2013). Several studies show synergetic effects when combining LRAs treatments (Darcis et al., 2015; Jiang et al., 2015; Laird et al., 2015; Martinez-Bonet et al., 2015). We, therefore, tested a combination of bryostatin-1 with either panobinostat or JQ1. Drugs were used at concentrations previously shown to be effective at reversing latency in other model systems (Archin, Liberty, et al., 2012; Bullen, Laird, Durand, Siliciano, & Siliciano, 2014; Laird et al., 2015; Spina et al., 2013).

To directly compare data from HIV_GKO_ infected cells with published *ex-vivo* results, we assessed LRA efficacy using PCR-based assays. We treated 5 million purified resting CD4^+^ T cells from four HIV infected individuals on suppressive ART (participant characteristics in Table 1) with single LRAs or combinations thereof, or vehicle alone for 24h. We then measured levels of intracellular HIV-1 RNA using primers and a probe that detect the 3' sequence common to all correctly terminated HIV-1 mRNAs (Shan et al., 2013). Of the LRAs tested individually, none were shown to have a statistically significant effect (n=4 - Figure 2A). Importantly, T-cell activation positive control, αCD3/CD28 (24.4-fold, Figure 2A), showed expected fold induction value (10 to 100-fold increases of HIV RNA in PBMCs (Bullen et al., 2014; Darcis et al., 2015; Laird et al., 2015)). When combining the PKC agonist bryostatin-1 with JQ1 or with panobinostat (fold-increases of 126.2- and 320.8-fold, respectively, Figure 2A), both combinations were highly more effective than bryostatin-1, JQ1 or panobinostat alone (fold-increases of 6.8-, 1.7- and 2.9-fold, respectively, Figure 3A), and even greater than the magnitude of induction stimulated by T-cell activation with αCD3/CD28. The synergetic relationship between those compounds was consistent with previous reports (Darcis et al., 2015; Jiang et al., 2015; Laird et al., 2015; Martinez-Bonet et al., 2015).

**Table 1.** Characteristics of HIV-1-infected study participants. ABC, abacavir; DRV, darunavir; FTC, emtricitabine; RPV, rilpivirine; RTV, ritonavir; TCV, tivicay; TDF, tenofovir; 3TC, lamivudine

| SCOPE ID | Age | Sex | Ethnicity | CD4 Count | Duration of Infection (years) | ART Regimen | Peak reported VL (copies ml<sup>-1</sup>) |
| --- | --- | --- | --- | --- | --- | --- | --- |
| 1597 | 56 | Male | Mixed | 469 | 19 | RPV/TDF/FTC | 45734 |
| 2147 | 59 | Male | Asian | 597 | 28 | RPV/TDF/FTC | 374000 |
| 2461 | 62 | Male | White | 664 | 32 | RPV, TCV | 20000 |
| 3162 | 54 | Male | White | 734 | 29 | RTV, DRV, ABC/TCV/3TC | 171000 |

**Fig. 2.**
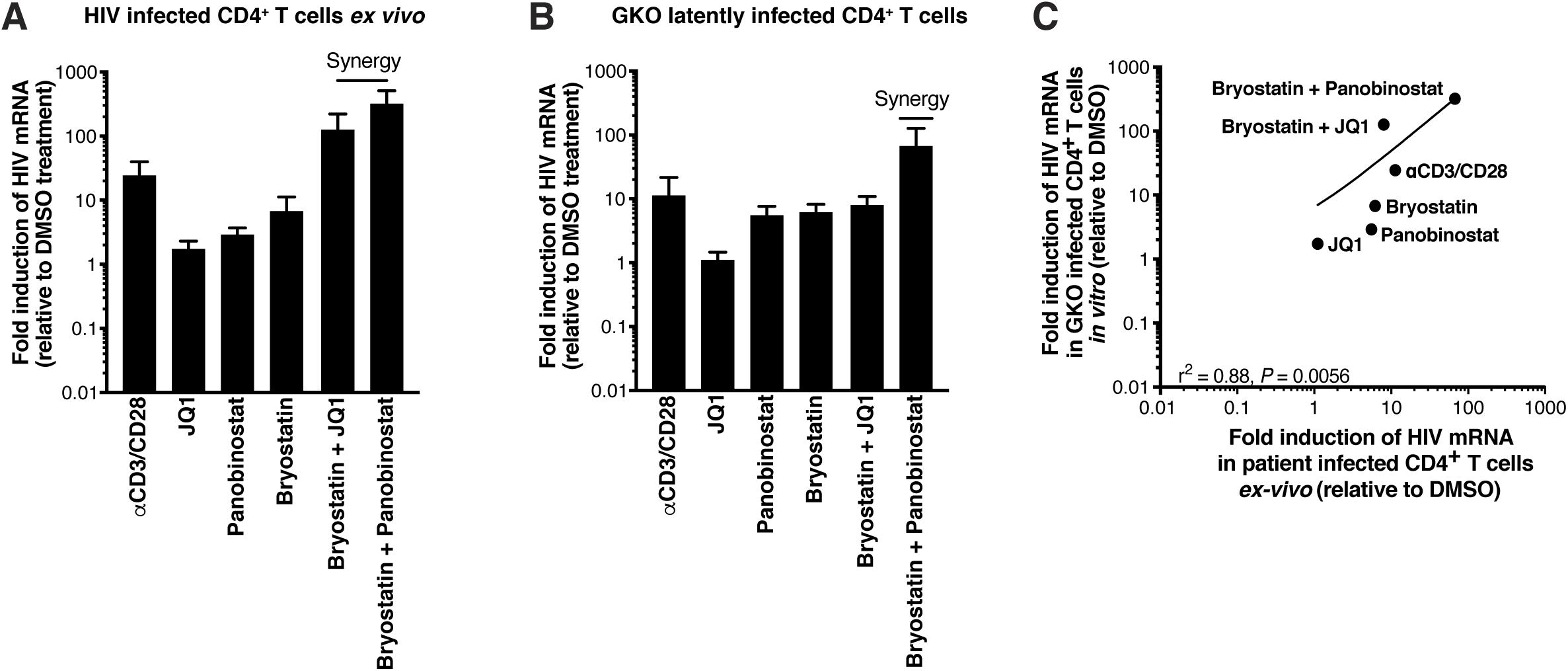
LRA efficacy in patient samples is predicted by activity in HIV_GKO_ latently infected cells. (A) Intracellular HIV-1 mRNA levels in rCD4s, obtained from infected individuals and treated *ex vivo* with a single LRA or a combination of two LRAs for 24-h, presented as fold induction relative to DMSO control. (n = 4, mean + SEM). (B) Intracellular HIV-1 mRNA levels in HIV_GKO_ latently infected CD4^+^ T-cells, and treated with a single LRA or a combination of two LRAs for 6h in presence of raltegravir, presented as fold induction relative to DMSO control. (n = 3, mean + SEM). (C) Correlation between intracellular HIV-1 mRNA levels quantified in either 6h stimulated HIV_GKO_ latently infected CD4^+^ T-cells, or 24h stimulated rCD4s from HIV infected patients, with a single LRA or a combination of two LRAs in presence of raltegravir.

**Fig. 3.**
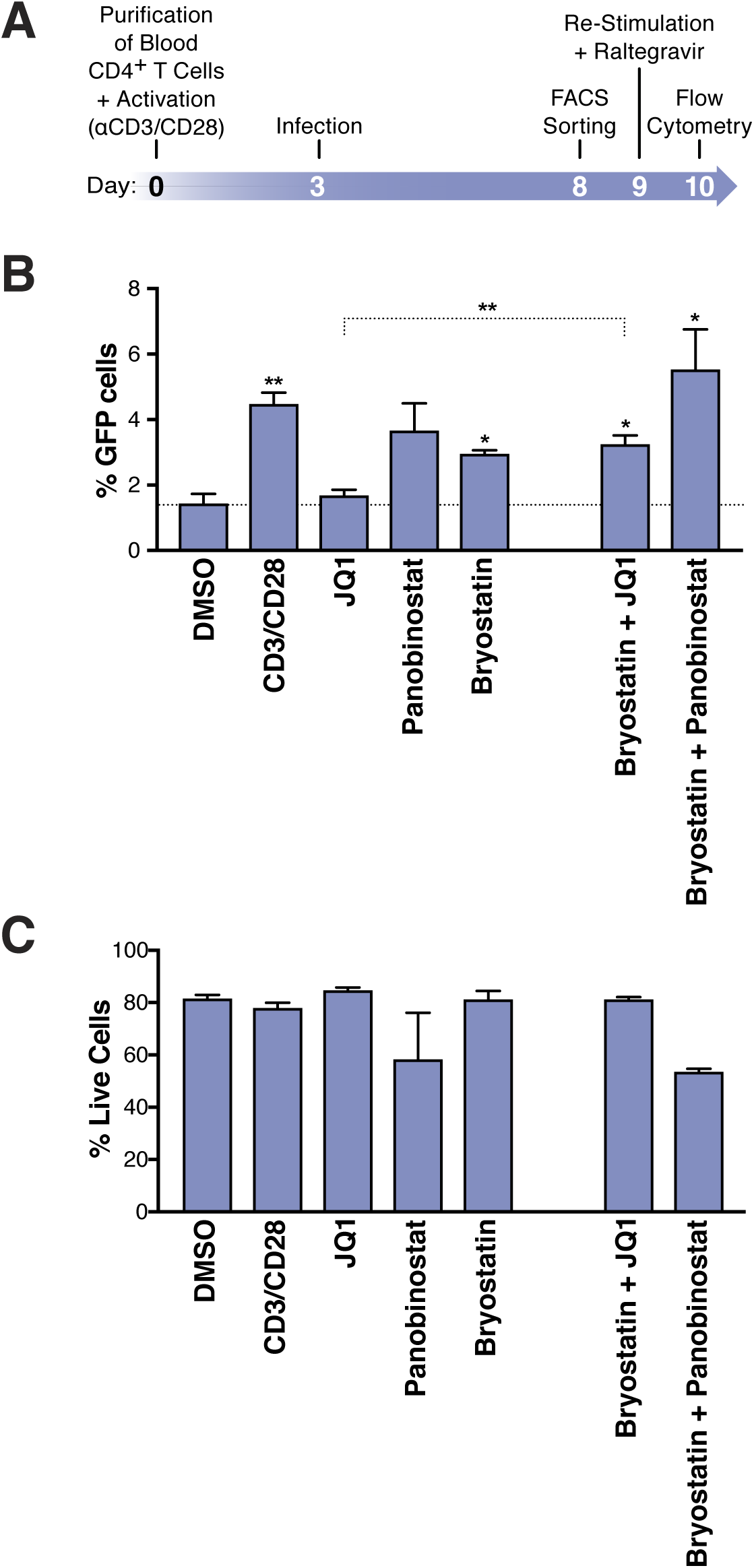
Few HIV_GKO_ latently infected primary CD4^+^ T cells are reactivated. (A) Schematic of experimental procedure with primary CD4^+^ T cells. Briefly, CD4^+^ T cells were purified from blood of healthy donors and activated for 72 h with αCD3/CD28 beads and 100 U/ml IL-2 before infection with HIV_GKO_. Five days post-infection, latently infected cells (csGFP-mKO2+) cells were sorted, put back in culture overnight and stimulated with different LRAs in presence of raltegravir for 24h before performing FACS analysis (B) Percentage of GFP+ cells are shown after stimulation of latently infected CD4^+^ T-cells with LRAs (n = 4, mean + SEM, paired t-test). (C) Histogram plot of percent live cells for each drug treatment (n = 3, mean + SEM, paired t-test). p-value: * p < 0.05, ** p < 0.01 relative to DMSO.

Measurement of intracellular HIV-1 mRNA in HIV_GKO_ latently infected cells showed an expected fold induction of latency in response to αCD3/CD28 (11.3-fold, Figure 2B). Second, JQ1, panobinostat, and bryostatin-1 alone all displayed mild to limited effect on reactivating HIV latency (fold-increases of 1.1-, 5.6- and 6.2-fold, respectively, Figure 2B). Finally, we observed low synergy when combining bryostatin and JQ1 (8-fold increase), but high synergy between bryostatin and panobinostat (67.3-fold increase).

All together, the data shown here are in agreement with current literature ((Darcis et al., 2015; Jiang et al., 2015; Laird et al., 2015; Martinez-Bonet et al., 2015)), and demonstrate that GKO virus *in vitro* closely mimics what is observed in patients’ samples (corelation rate r^2^=0.88, p=0.0056 - Figure 2C).

### HIV-1 LRAs target a minority of latently infected primary CD4^+^ T cells

Current assays have evaluated the efficacy of different LRAs by relative quantification of the impact of LRAs on the latent reservoir by measuring viral RNA, DNA or proteins (Figure 2A). The use of dual-fluorescent HIV reporters, however, provides a tool to quantify directly the fraction of cells in different states.

To quantify the absolute proportion of induced latently infected cells following LRAs treatment, primary CD4^+^ T cells were infected with HIV_GKO_, and cultured for 5 days (in presence of IL-2) before sorting the different populations. Cells were allowed to rest overnight and then treated for 24h with the various LRAs (same drugs concentrations as in Figure 2) (Figure 3A). Culture of DMSO-treated latently infected primary CD4^+^ T cells produced little spontaneous reactivation (average of four experiments: 1.4% of GFP+ cells). Of all LRAs, neither JQ1 (1.7%) nor panobinostat (3.7%) significantly reactivated latently infected cells, even though the mean reactivation potential of panobinostat was twofold higher than that of JQ1 (Figure 3B). Panobinostat demonstrated toxicity in primary cells (Figure 3C). Treatment of latent CD4^+^ T cells with bryostatin-1 (3%) led to significant and similar fold reactivation of the latent population, but not as strong as the positive control αCD3/CD28 (4.5%).

However, we did not observe synergy, nor a partial additive effect when combining byrostatin-1 and JQ1 (3.3%. vs bryostatin or JQ1 alone: 3%, and 1.7%, respectively). When treating cells with both bryostatin-1 and panobinostat (5.5%), we visualized a partial additive effect (Figure 3B).

All together, these data show that LRAs reactivate only a very small proportion of latent cells in primary CD4^+^ T cells. Surprinsingly, the positive control αCD3/CD28, described in the literature as being the most potent reactivation, only reactivated up 3.1% (when substracting spontaneous reactivation observed in DMSO treated samples) of latently infected cells. The maximal induction of the whole latent population was achieved by combining bryostatin + panobinostat with 4.1% of reactivation (when substracting spontaneous reactivation observed in DMSO treated samples).

### Small fractional rate of latency reactivation is not explained by low cellular response to activation signals

Our data showed that fold inductions of HIV_GKO_ latently infected cells using different LRAs were in agreement with current literature ((Jiang et al., 2015; Mehla et al., 2010; Mitchell et al., 2004; Whitney et al., 2014)) (Figure 2B), however when looking at the absolute number of cells being reactivated, we found a surprisingly low fraction of reactivated latently infected cells in primary CD4^+^ T cells in response to all agents (Figure 3B). This was particularly surprising in response to αCD3/CD28 stimulation, as current models for HIV latency point that the state of T cells activation dictates the transcriptional state of the virus. Treatment of latently infected cells with αCD3/CD28 stimulated HIV production in only approximately 5% of the cells while the other 95% remained latent, even though > 95% of the cells were expressing T-cell activation-associated surface markers CD69 and CD25 (Figure S1).

To rule out the possibility that non-reactivated latently infected cells (NRLIC) failed to reactivate the provirus due to inefficient response to T-cell activation signals, we analyzed T-cell activation markers within the different populations (i.e., within uninfected, NRLIC and reactivated latently infected cells (RLIC); Figure 4). Briefly, 72h stimulated-CD4^+^ T cells were infected with HIV_GKO_; 4 days post-infection, GFP-cells were sorted, and allowed to rest overnight before stimulating the cells with αCD3/CD28. Cells were stained for CD25 and CD69 expression 24h later. Both RLIC and NRLIC were mainly composed of strongly activated cells (77.3% and 63.9% respectively; CD25+/CD69+) and very few (< 3%) non-stimulated cells (CD25-/CD69-) (Figure 4). The RLIC population comprised a small but statistically significant increase of cells expressing only the early T-cell activation marker (CD25-/CD69+), as well as a non-significant decrease of mature stimulated T cells. Overall, the comparison of the latent populations showed very little difference.

**Fig. 4.**
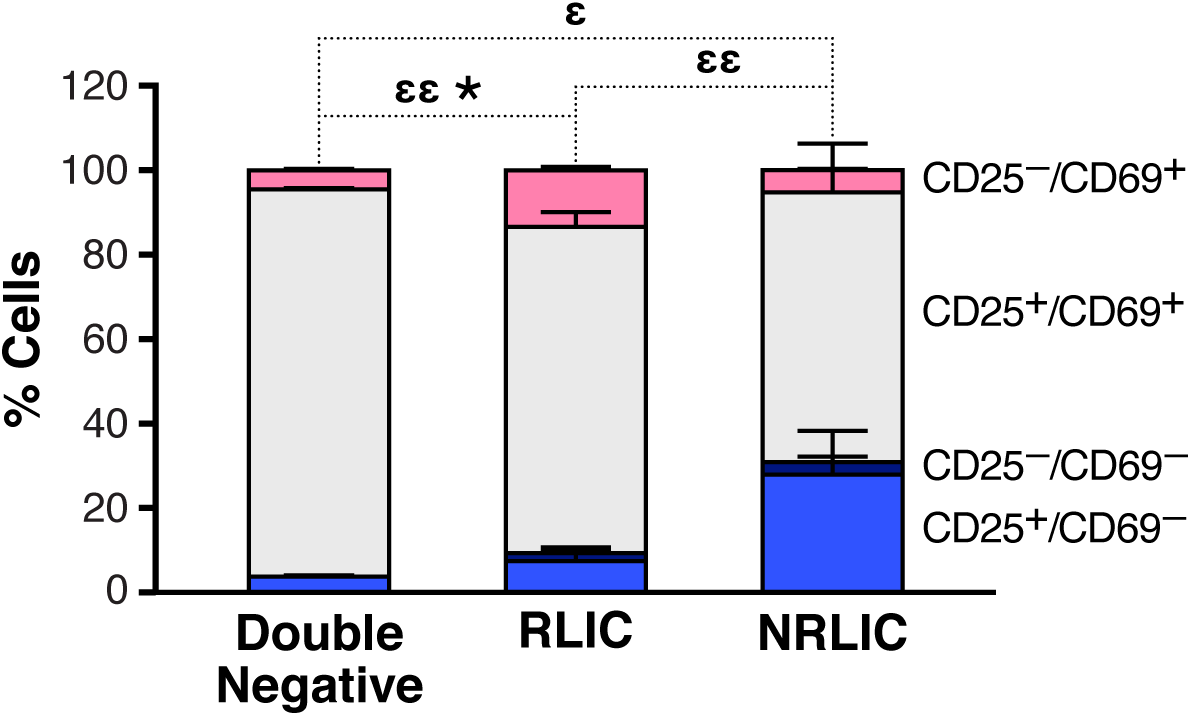
Low-level latency reactivation is not explained by low cellular responses to activation signals. T-cell activation patterns between reactivated and non-reactivated latently infected cells. Briefly, CD4^+^ T-cells were purified from blood of four healthy donors and activated for 72 h with αCD3/CD28 beads and 20 U/ml IL-2 before infection with HIV_GKO_. At 4 days post-infection, csGFP-were sorted, cultured overnight and stimulated with αCD3/CD28 in presence of raltegravir. At 24 h post-treatment, cells were stained for CD25 and CD69 activation markers before performing FACS analysis. p-value: *p<0.05 for CD25+/CD69+ population, and ε p < 0.05, εε p < 0.01 for CD25-/CD69+ population.

In summary, after validating the efficacy of the different LRAs used in this study, and the accuracy of our dual-fluorescent reporter, we demonstrate that, in the HIV_GKO_ latency primary CD4^+^ T cell model, only a small fraction of latently infected cells undergoes viral reactivation in response to different activation signals, even though most of the cells are targeted and respond to effective stimuli.

### Integration sites and host transcriptional activities affect the fate of HIV-1 infection

The latent reservoir has been difficult to characterize, and whether or not the genomic location of the integration affects latency is debated (Chen et al., 2016; Dahabieh et al., 2014; Jordan et al., 2003; Jordan et al., 2001; Sherrill-Mix et al., 2013).

To determine whether the site of integration modulated the reactivation of latent HIV, primary CD4^+^ T-cells were infected with HIV_GKO_ 3 days post-activation. At 5 days post-infection, productively infected cells (GFP+, PIC) were sorted and frozen, and the GFP-populations (latent and uninfected) were isolated and treated with αCD3/CD28. 48h post-induction, the NRLIC and RLIC populations were isolated. Nine libraries (three donors, three samples/donor: PIC, RLIC, NRLIC) were constructed from genomic DNA as described (Cohn et al., 2015) and analyzed by high-throughput sequencing to locate the HIV provirus within the human genome. A total of 1,803 virus integration sites were determined: 960 integrations in PIC, 681 in NRLIC, and 162 in RLIC.

First, we explored whether integration within genes involved in T-cell activation predicted infection reactivation fate. To do so, we compared our HIV integration dataset with a previously published dataset that profiled gene expression from resting and activated (48h - αCD3/CD28) CD4^+^ T cells from PBMCs of healthy individuals (Ye et al., 2014). The analysis revealed that most of the αCD3/CD28-induced latent proviruses were not integrated in genes responsive to T-cell activation signals (Figures 5A and 5B). Notably, PIC and RLIC integration events clearly targeted genes whose basal expression was significantly higher than genes targeted in NRLIC, both in activated and resting T cells (Figure 5C).

**Fig. 5.**
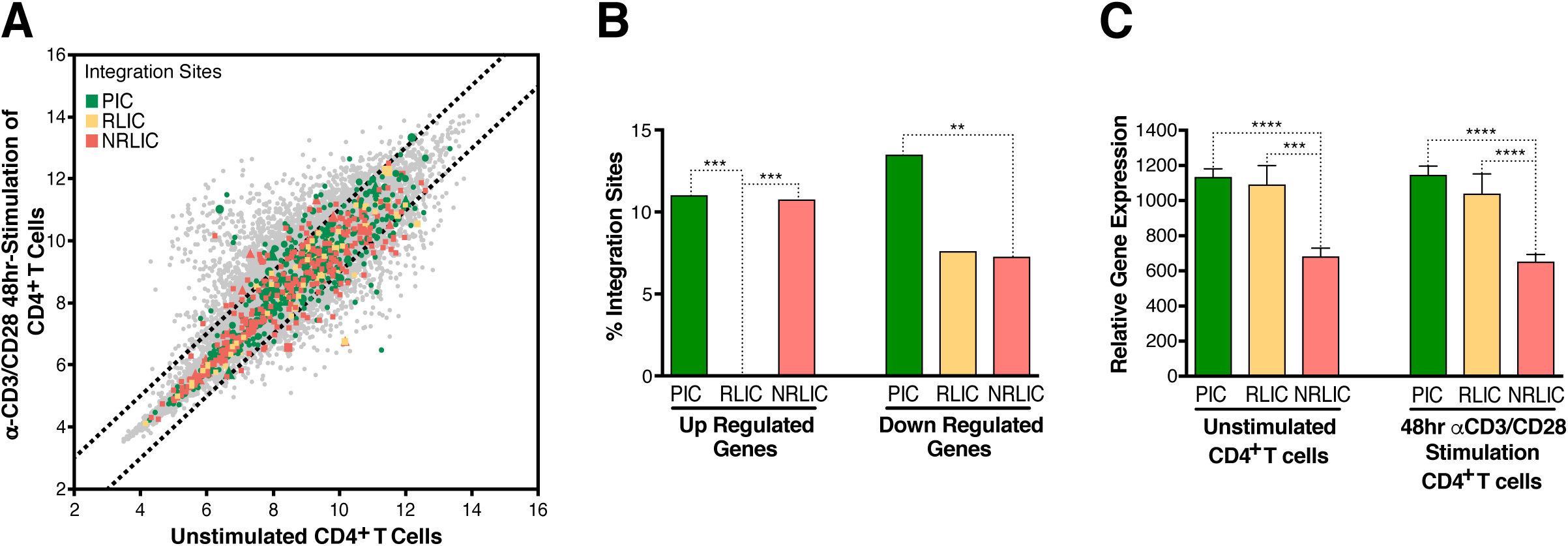
Relative expression of HIV-1 integration targeted genes for each population, before or after TCR activation. (A) Scatter charts showing primary CD4^+^ T-cell gene expression changes after 48h of stimulation with αCD3/CD28 beads. Integration sites displayed outside of the two solid gray lines were targeted genes whose expression is at least +/-twofold differentially expressed after 48h stimulation. Plot points size can be different, the bigger the plot point is, the more integration events happened within the same gene. (B) Fraction of integration sites from the different populations PIC, RLIC or NRLIC, integrated within genes whose expression is at least +/-twofold differentially expressed after 48h of αCD3/CD28 stimulation (**p < 0.01; ***p < 0.001; two-proportion z test). (C) Relative expression of genes targeted by HIV-1 integration in PIC, RLIC or NRLIC before TCR stimulation and after αCD3/CD28 stimulation (n=3, mean + SEM, paired t-test). ***p < 0.001; ****p < 0.0001

Secondly, we investigated whether different genomic regions were associated with productive, inducible or non-inducible latent HIV-1 infection. In agreement with previous studies (Cohn et al., 2015; Dahabieh et al., 2014; Maldarelli et al., 2014; Wagner et al., 2014), the majority of integration sites were found within genes in each population (Figure 6A), although the proportion of genic integrations in NRLIC was significantly lower than in PIC and RLIC samples. Moreover, integration events in the PIC and RLIC populations were more frequent in expressed regions (sum of low + medium + high expressed genes = 64% and 58%, respectively), while these regions were significantly less represented in the NRLIC (31%) (Figure 6B). In addition, genic integration events were more frequent in the introns for each population (> 65%, Figure 6C). Finally, viral orientation of the provirus did not correlate with the fate of HIV infection or the reversal of HIV latency (Figure 6D).

**Fig. 6.**
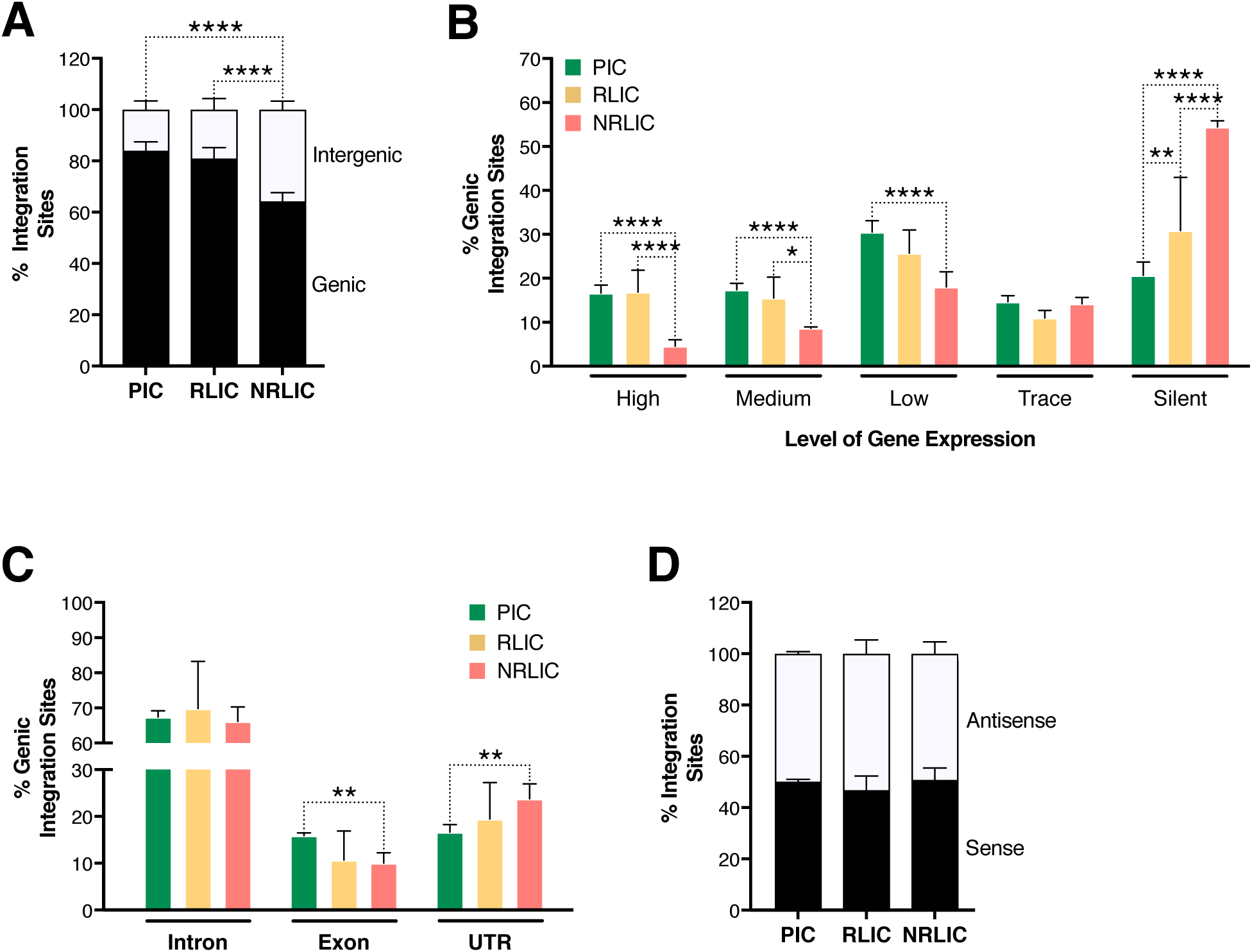
Insertion landscapes of HIV-1. (A) Proportion of mapped insertions that are in genic or intergenic regions. (B) Proportion of integrations in transcribed regions with high (top 1/8), medium (top 1/4–1/8), low expression (top 1/2–1/4), trace (bottom 1/2) or silent (0) expression. p-value: (C) Proportion of unique genic integrations located in introns, exons, UTR or promoters. (D) Transcriptional orientation of integrated HIV-1 relative to host gene. *p < 0.05; **p < 0.01; ***p < 0.001; ****p < 0.0001 using two-proportion z test.

### Integration sites and chromatin context affect the fate of HIV-1 infection

Chromatin marks, such as histone post-translational modifications (e.g., methylation and acetylation) and DNA methylation, are involved in establishing and maintaining HIV-1 latency (De Crignis & Mahmoudi, 2017). We examined 500 bp regions centered on all integration sites in each population for several chromatin marks by comparing our data with several histone modifications and DNaseI ENCODE datasets. We first looked at distinct and predictive chromatin signatures, such as H3K4me1 (active enhancers), H3K36m3 (associated with active transcribed regions), H3K9m3 and H3K27m3 (repressive marks of transcription) (reviewed in (Kumar, Darcis, Van Lint, & Herbein, 2015; Shlyueva, Stampfel, & Stark, 2014)). All three populations had distinct profiles, although productive and inducible latent infection profiles appeared most similar (Figure 7A). The analysis showed that PIC integrated in active chromatin (i.e., transcribed genes - H3K36me3 or enhancers - H3K4me1), while NRLIC integration appeared biased toward heterochromatin (H3K27me3 and H3K9me3) and non-accessible regions (DNase hyposensitivity). Interestingly, the analysis also showed that RLIC population shared features with PIC regarding H3K36me3, H3K4me1, and H3K27me3 marks, but also with NRLIC regarding H3K9me3 mark and DNase accessibility.

**Fig. 7.**
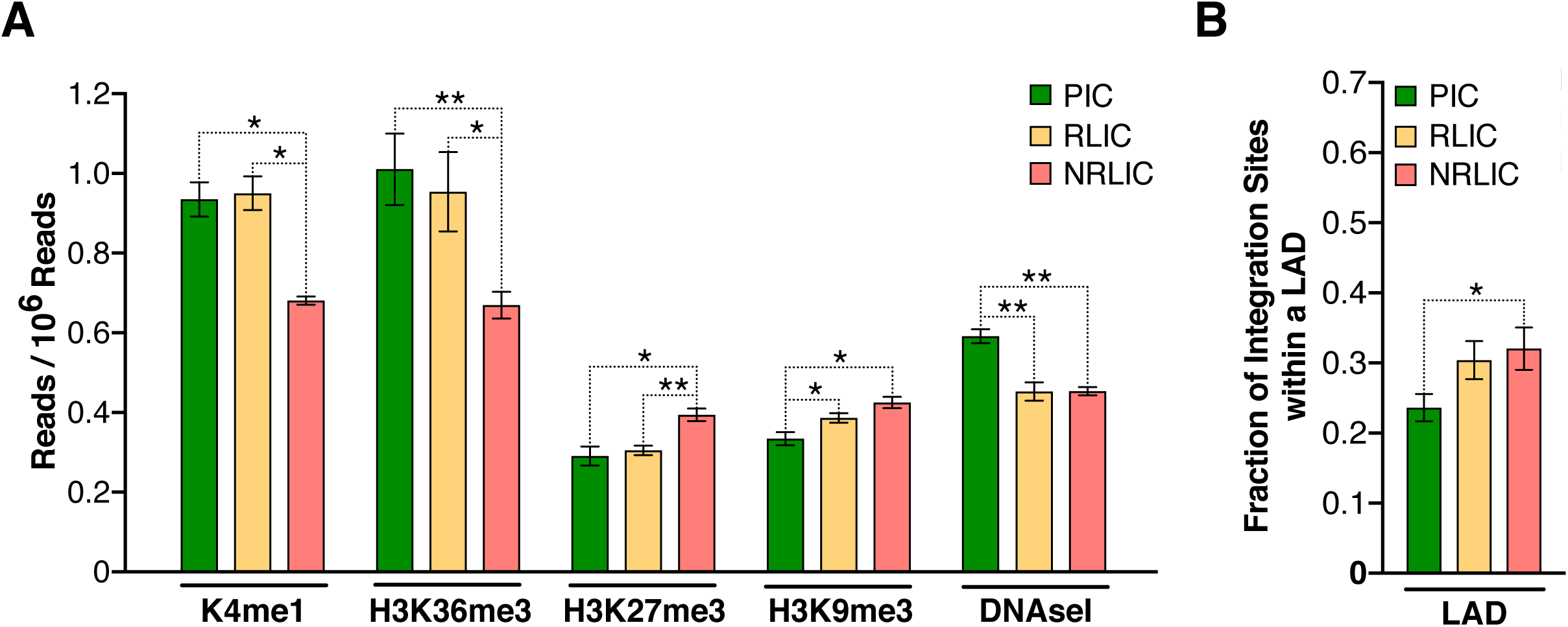
Epigenetics marks and nuclear localization of HIV-1 integration sites. (A) 500 bp centered on HIV-1 integration sites for each population were analyzed for the presence of H3K4me1 (active enhancers), H3K36m3 (active transcribed regions), H3K9m3 and H3K27m3 (repressive marks of transcription), and DNA accessibility (DNAseI). (B) Nuclear localization of HIV-1 integration sites. Quantification was based on inside a LAD (=1) or outside (=0), which means the Y axis represents the fraction of integrations within a LAD. (n = 3–4 ENCODE donors, mean + SEM, paired t-test). *p < 0.05; **p < 0.01; ***p < 0.001)

In a related study, Marini *et al.* show that HIV-1 mainly integrates at the nuclear periphery (Marini et al., 2015). We examined the topological distribution of integration sites from each population inside the nucleus by analyzing our data using a previously published dataset of lamin-associated domains (LADs) (Guelen et al., 2008). LADs consist of H3K9me2 heterochromatin and are present at the nuclear periphery. Analysis showed that integration sites from NRLIC were in LADs to a significantly higher degree (32%) than productive integrations (23.6%) (p < 0.05, Figure 7B). Integration sites from RLIC also tended to be integrated in LAD (30.4%).

Overall, these data show similar features between productively infected and inducible latently infected cells, while non-reactivated latently infected cells appear distinct from the other populations. These findings indicate a prominent role of the site of integration and the chromatin context for the fate of the infection itself, as well as for latency reversal.

## DISCUSSION

Dual-color HIV-1 reporters are unique and powerful tools (Calvanese et al., 2013; Dahabieh, Ooms, Simon, & Sadowski, 2013), that allow for the identification and the isolation of early latently infected cells from productively infected cells and uninfected cells. Latency is established very early in the course of HIV-1 infection (Archin, Vaidya, et al., 2012; Chun et al., 1998; Whitney et al., 2014) and, until the advent of dual-reporter constructs, no primary HIV-1 latency models have allowed the study of latency heterogeneity at this very early stage. Importantly, the comparison of data obtained from distinct primary HIV-1 latency models is complicated as some models are better suited to detect latency establishment (e.g., dual-reporters), while others are biased towards latency maintenance (e.g., Bcl2-transduced CD4^+^ T cells). The use of env-defective viruses limits HIV replication to a single-round and, thereby limits the appearance of defective viruses (Bruner et al., 2016). Thus, theoretically, most of the HIV_GKO_ latent provirus can be induced to produce infective particles, although several rounds of activation may be needed using differently acting LRAs.

In this study, we describe and validate an improved version of HIV_DuoFluoi_, previously developed in our laboratory (Calvanese et al., 2013), which accurately allows for: a) the quantification of latently infected cells, b) the purification of latently infected cells, and c) the evaluation of the “shock and kill” strategy, since HIV_GKO_ recapitulates LRAs response observed with HIV infected cells from patients. The motivation of the study is to better understand the mechanisms of HIV reactivation in primary cells since none of the interventions conducted thus far in patients has reduced the size of the latent HIV-1 reservoir *in vivo* (Rasmussen & Lewin, 2016). Our data highlight two important facts: a) cell-associated HIV RNA quantification does not reflect the number of cell undergoing viral reactivation, and b) a small portion of the latent proviruses (< 5%) is reactivated, although LRAs target the whole latent population. Hence, even if cells harboring reactivated virus die, this small reduction would likely remain undetectable when quantifying the latent reservoir *in vivo.* Our data are in agreement with previous reports, which show that levels of cellular HIV RNA and virion production are not correlated, and that the absolute number of cells being reactivated by αCD3/CD28 is indeed limited to a small fraction of latently infected cells (Cillo et al., 2014; Sanyal et al., 2017; Yucha et al., 2017). Using our dual-fluorescence reporter, we confirme these findings, and extend these observations to LRAs combination. However, although LRAs combination shows synergy when measuring cell-associated HIV RNA, we do not find such synergy, but only partial addive effect, when quantifying the absolute number of induced latently infected cells. Our work, and other’s (Cillo et al., 2014; Sanyal et al., 2017; Yucha et al., 2017), demonstrate the importance of single cell analysis when it comes to the evaluation of potential LRAs. Indeed, it is necessary to determine wheter potential increases in HIV RNA after stimulation in a bulk population result from a small number of highly productive cells, or from a larger but less productive population, as these two outcomes likely have very different impact on the latent reservoir.

Our data further highlight the heterogeneous nature of the latent reservoir (Chen et al., 2016; Ho et al., 2013). Active infections lead to the production of virions, whereas latent infections lead to transcriptional repression. There is currently a limited understanding as to why some latently infected cells are capable of being induced, while others are not. It is possible that transcriptional repression is influenced by the integration site context, which would affect both viral transcription and HIV-1 latency reversal (Chen et al., 2016). Since HIV_GKO_ allows the isolation of productively infected cells and reactivated latent cells from those that do not reactivate, it provides a unique opportunity to explore the impact of HIV integration on the fate of the infection and on the ability of different latent HIV to become reactivated.

Different integration site-specific factors contribute to latency, such as the chromatin structure of the HIV-1 provirus, including adjacent loci but also the provirus location in the nucleus (Lusic & Giacca, 2015; Lusic et al., 2013). Viral integration is a semi-random process (Bushman et al., 2005) in which HIV-1 preferentially integrates into active genes (Barr et al., 2006; Bushman et al., 2005; Demeulemeester, De Rijck, Gijsbers, & Debyser, 2015; Ferris et al., 2010; Han et al., 2004; Lewinski et al., 2006; Mitchell et al., 2004; Schroder et al., 2002; Sowd et al., 2016; Wang, Ciuffi, Leipzig, Berry, & Bushman, 2007). LEDGF, one of the main chromatin-tethering factors of HIV-1, binds to the viral integrase and to H3K36me3, and to a lesser extent to H3K4me1, thus directing the integration of HIV-1 into transcriptional units (Daugaard et al., 2012; Eidahl et al., 2013; Pradeepa, Sutherland, Ule, Grimes, & Bickmore, 2012). Also CPSF6, which binds to the viral capsid, markedly influences integration into transcriptionally active genes and regions of euchromatin (Sowd et al., 2016), explaining how HIV-1 maintains its integration in the euchromatin regions of the genome independently of LEDGF (Quercioli et al., 2016). Several studies have characterized the integration sites, however, these analyses have been restricted to productive infections.

Consistent with previous results and using ENCODE reference datasets, our data show that HIV-1 preferentially integrates into genic regions. Productive proviruses predominantly target actively transcribed regions, as predicted by chromatin signatures, such as H3K36m3, found in stably transcribed genes (Marini et al., 2015; Wang et al., 2007) and H3K4me1 (Chen et al., 2016), marking active enhancers. On the other hand, non-inducible latent proviruses are observed to be integrated into silenced chromatin, with low DNaseI accessibility and marked by H3K27me3. Although HIV-1 preferentially integrates into the peripheral nuclear compartment (Albanese, Arosio, Terreni, & Cereseto, 2008; Burdick, Hu, & Pathak, 2013; Marini et al., 2015; Quercioli et al., 2016), integration is normally strongly disfavored in the heterochromatic condensed regions in lamin-associated domains (LADs). Here, when using a previously published dataset of LADs (Guelen et al., 2008; Marini et al., 2015), we show that HIV integration does occur in LADs, but that it results in a latent provirus with low probability to be reactivated.

Taking into account the preferential HIV-1 integration into open chromatin regions, it remains to be determined how the heterochromatic status of the provirus is established during latency. For latency to occur, viruses initially integrated into permissive regions of the genome may become repressed (i.e., during the transition of the target cell towards a quiescent state). During this transition, repressive chromatin marks are deposited on the site of integration, CpG islands within promoters are methylated, and transcription factors are depleted (Blazkova et al., 2009; du Chene et al., 2007; Friedman et al., 2011; Imai, Togami, & Okamoto, 2010; Kauder, Bosque, Lindqvist, Planelles, & Verdin, 2009; Marcello et al., 2003; Sabo, Lusic, Cereseto, & Giacca, 2008; Williams, Kwon, Chen, & Greene, 2007). Another possibility is that the virus is integrated directly into heterochromatic regions, which subsequently spread to silence the viral genome. Our data suggest that both scenarios occur in cells, but result in different fates of the infection. Indeed, although HIV is preferentially integrated in open chromatin, especially for the productive population, a substantial fraction of integrations occurs in heterochromatin (Jordan et al., 2003). This population is resistant to viral reactivation concomitant with T-cell activation.

Importantly, we identify a unique rare population among the latent cells that can be reactivated. In contrast to the non-inducible latent infections, the latency reversal of inducible latent proviruses might be explained by integration in an open chromatin context, similar to integration sites for productive proviruses, followed by subsequent heterochromatin formation and proviral silencing. As a consequence, the distinct genomic profiles between induced and non-induced latent provirus opens up new possibilities for cure strategies. Indeed, the “shock and kill” strategy aims to reactivate and eliminate every single replication-competent latent provirus, since a single remaining cell carrying a latent inducible provirus could, in theory, reseed the infection. However, our study, and others’, point out several complications to the “shock and kill” strategy. First, LRAs only reactivate a limited fraction of latent proviruses and, within the latent population, a large fraction is resistant to reactivation. It is likely that some of the non-induced proviruses will reactivate after several rounds of activation, due to the stochastic nature of HIV activation (Dar et al., 2012; Ho et al., 2013; Singh et al., 2010; Weinberger et al., 2005). However, our data show that the provirus is efficiently silenced by cellular mechanisms. As such, reactivation of these dormant proviruses would likely require intense effort and more potent LRAs (Rouzine, Razooky, & Weinberger, 2014). Second, the cells harboring the induced latent proviruses are not immediately killed, implying that immunomodulatory approaches, in addition of more potent LRAs, are likely required to achieve a cure for HIV infection (Shan et al., 2012).

In conclusion, in addition to eliminating cells with productive and inducible proviruses, it might be relevant to explore other strategies to a functional HIV-1 cure to deal with the remaining latent reservoir. The “block and lock” approach would consist of enforcing the cellular mechanisms to maintain latent provirus silenced (Besnard et al., 2016; Vranckx et al., 2016). For a functional cure, a stably silenced, non-reactivatable provirus is tolerated.

## MATERIALS AND METHODS

### Patients’ samples

Four HIV-1-infected individuals, who met the criteria of suppressive ART, undetectable plasma HIV-1 RNA levels (<50 copies/ml) for a minimum of six months, and with CD4^+^ T cell count of at least 350 cells/mm^3^, were enrolled. The participants were recruited from the SCOPE cohort at the University of California, San Francisco. Table 1 details the characteristics of the study participants.

### Plasmids construction

To construct HIV_GKO_, the csGFP sequence was designed and ordered from Life Technologies. The sequence was cut out from Life Technologies’ plasmid with BamHI and XhoI and cloned into DuoFluo, previously cut with the same enzymes (DuoFluo-csGFP). HIV_GKO_ was creating by PCR overlapping: csGFP-EF1α (Product 1) was PCR amplified from DuoFluo using primers P1: 5' for-GATTAGTGAACGGATCCTTGGCAC-3' and P2: 5’ rev-GGCTTGATCACAGAAACCATGGTGGCGACCGGTAGCGC-3'. mKO2 (Product 2) was PCR amplified from Brian Webster’s plasmid (kind gift from Warner Greene) using primers P3: 5' for-GCGCTACCGGTCGCCACCATGGTTTCTGTGATCAAGCC-3’ and P4: 5’ rev-CTCCATGTTTTTCCAGGTCTCGAGCCTAGCTGTAGTGGGCCACGGC-3’. Finally, we amplified the 3’LTR sequence (Product 3) from RGH plasmid (Dahabieh et al., 2013) using primers P5: 5’ for-GCTCGAGACCTGGAAAAACATGGAG-3’ and P6: 5’ rev-GTGCCACCTGACGTCTAAGAAACC-3’, to add a fragment containing the AatII restriction site, in order to ligate the csGFP-EF1α-mKO2 cassette into pLAI(Peden, Emerman, & Montagnier, 1991). We then did sequential PCRs: products 1 and 2 were amplified using primers P1 and P4. PCR product (1+2) was mixed with product 3 and PCR amplified with P1 and P6 thus creating the full cassette. The cassette csGFP-EF1α-mKO2 was then digested with BamHI and AatII, and cloned into pLAI previously digested with the same enzymes to create HIV_GKO_.

Of note, the Envelope open reading frame was disrupted by the introduction of a frame shift at position 7136 by digestion with KpnI, blunting, and re-ligation.

To construct GKO-∆U3 3’LTR, we cloned a ∆U3 linker from pTY-EFeGFP (Chang, Urlacher, Iwakuma, Cui, & Zucali, 1999; Cui, Iwakuma, & Chang, 1999; Iwakuma, Cui, & Chang, 1999; Zolotukhin, Potter, Hauswirth, Guy, & Muzyczka, 1996) into the KpnI/SacI sites of the 3’ LTR in HIV_GKO_.

### Virus production

Pseudotyped HIV_DuoFluoi_, HIV_GKO_, and HIV R7/E-/GFP(Jordan et al., 2003) viral stocks were generated by co-transfecting (standard calcium phosphate transfection method) HEK293T cells with a plasmid encoding HIV_DuoFluoi_, HIV_GKO_ or R7/E^-^/GFP, and a plasmid encoding HIV-1 dual-tropic envelope (pSVIII-92HT593.1) or the envelope G glycoprotein of the vesicular stomatitis virus (VSV-G for Jurkat infections). Medium was changed 6–8 h post-transfection, and supernatants were collected after 48 h, centrifuged (20 min, 2000 rpm, RT), filtered through a 0.45 μM membrane to clear cell debris, and then concentrated by ultracentrifugation (22,000g, 2 h, 4°C). Concentrated virions were resuspended in complete media and stored at −80°C. Virus concentration was estimated by p24 titration using the FLAQ assay (Marianne Gesner, 2014).

### Primary cell isolation and cell culture

CD4^+^ T cells were extracted from peripheral blood mononuclear cells (PBMCs) from continuous-flow centrifugation leukophoresis product using density centrifugation on a Ficoll-Paque gradient (GE Healthcare Life Sciences). Resting CD4^+^ lymphocytes were enriched by negative depletion with an EasySepHuman CD4^+^ T Cell Isolation Kit (Stemcell). Cells were cultured in RPMI medium supplemented with 10% fetal bovine serum, penicillin/streptomycin and 5 μM saquinavir.

Primary CD4^+^ T cells were purified from healthy donor blood (Blood Centers of the Pacific, San Francisco, CA, and Stanford Blood Center), by negative selection using the RosetteSep Human CD4^+^ T Cell Enrichment Cocktail (StemCell Technologies). Purified resting CD4^+^ T cells from HIV-1 or healthy individuals were cultured in RPMI 1640 medium supplemented with 10% FBS, L-glutamine (2 mM), penicillin (50 U/ml), streptomycin (50 mg/ml), and IL-2 (20 to 100 U/ml) (37°C, 5% CO**2**). Spin-infected primary CD4^+^ T cells were maintained in 50% of complete RPMI media supplemented with IL-2 (20–100 U/ml) and 50% of supernatant from H80 cultures (previously filtered to remove cells) without beads. Medium was replenished every 2 days until further experiment.

HEK293T cells were obtained from ATCC. Feeder cells H80 was a kind gift from Jonathan Karn. H80 cells were cultured in RPMI 1640 medium supplemented with 10% fetal bovine serum (FBS), L-glutamine (2 mM), penicillin (50 U/ml), and streptomycin (50 mg/ml) (37°C, 5% CO**2**). HEK293T cells were cultured in DMEM medium supplemented with 10% FBS, 50 U/ml penicillin, and 50 mg/ml streptomycin.

### Cell infection

Purified CD4^+^ T cells isolated from healthy peripheral blood were stimulated with αCD3/CD28 activating beads (Life Technologies) at a concentration of 0.5 bead/cell in the presence of 20–100 U/ml IL-2 (PeproTech) for three days. All cells were spinoculated with either HIV_DuoFluoi_, HIV_GKO_ or HIV D3U-GKO at a concentration of 300 ng of p24 per 1.10^6^ cells for 2 h at 2000rpm at 37°C without activation beads.

Infected cells were either analyzed by flow cytometry or sorted 4–5 days post-infection.

### Latency-reversing agent treatment conditions

CD4^+^ T cells were stimulated for 24h unless stipulated differently, with latency-reversing agents at the following concentrations for all single and combination treatments: 10 nM bryostatin-1, 1 μM JQ1, 30 nM panobinostat, αCD3/CD28 activating beads (1 bead/cell), or media alone plus 0.1% (v/v) DMSO. For all single and combination treatments, 30 μM Raltregravir (National AIDS Reagent Program) was added to media. Concentrations were chosen based on Laird *et al.* paper (Laird et al., 2015).

### Staining, flow cytometry and cell sorting

Cells from Figure 4 were stained with α-CD69-PE-Cy7 and α-CD25-APC (BD Bioscience) and fixed in 2% paraformaldehyde.

Before collecting data using the FACS LSRII (BD Biosciences), cells were stained with violet Live/Dead Fixable Dead Cell Stain (Life Technologies) and fixed with 2% formaldehyde. Analyses were performed with FlowJo V10.1 software (TreeStar).

Sorting of infected CD4^+^ T cells was performed with a FACS AriaII (BD Biosciences) based on their GFP and mKO2 fluorescence marker at 4–5 days post-infection, and placed back in culture for further experimentation.

### DNA, RNA and protein extraction, qPCR and western blot

RNA and proteins (Figures 1B and 1C) were extracted with PARIS™ kit (Ambion) according to manufacturer’s protocol from same samples. RNA was retro-transcribed using random primers with the SuperScript II Reverse Transcriptase (Invitrogen) and qPCR was performed in the AB7900HT Fast Real-Time PCR System, using 2X HoTaq Real Time PCR kit (McLab) and the appropriate primer-probe combinations described in (Calvanese et al., 2013). Quantification for each qPCR reaction was assessed by the ddCt algorithm, relative to Taq Man assay GAPDH Hs99999905_m1. Protein content was determined using the Bradford assay (Bio-Rad) and 20 μg were separated by electrophoresis into 12% SDS-PAGE gels. Bands were detected by chemiluminescence (ECL Hyperfilm Amersham) with anti-Vif, HIV-p24 and a-actin (Sigma) primary antibodies.

Total RNA (Figures 3A and 3B) were extracted using the Allprep DNA/RNA/miRNA Universal Kit (Qiagen) with on-column DNAase treatment (Qiagen RNase-Free DNase Set). Cellular HIV mRNA levels were quantified with a qPCR TaqMan assay using primers and probes as described (Bullen et al., 2014) on a ViiA 7 Real-Time PCR System (Life Technologies). Cell-associated HIV mRNA copy numbers were determined in a reaction volume of 20 *μ*L with 10 *μ*L of 2x TaqMan^®^ RNA to Ct™ 1 Step kit (Life Technologies), 4 pmol of each primer, 4 pmol of probe, 0.5 *μ*L reverse transcriptase, and 5 *μ*L of RNA. Cycling conditions were 48°C for 20 min, 95°C for 10 min, then 60 cycles of 95°C for 15 sec and 60°C for 1 min. Real-time PCR was performed in triplicate reaction wells, and cell-associated HIV mRNA was normalized to cell equivalents using human genomic GAPDH expression by qPCR and applying the comparative Ct method (Vandesompele et al., 2002).

### HIV integration site libraries and computational analysis

HIV integration site libraries and computational analysis were executed in collaboration with Lilian B. Cohn and Israel Tojal Da Silva as described in their published paper (Cohn et al., 2015), with a few small changes added to the computational analysis pipeline. First, we included integration sites with only a precise junction to the host genome. Second, to eliminate any possibility of PCR mispriming, we have excluded integration sites identified within 100bp (50bp upstream and 50bp downstream) of a 9bp motif identified in our LTR1 primer: TGCCTTGAG. Thirdly we have merged integration sites within 250bp and have counted each integration site as a unique event. The list of integration sites for each donor and each population can be found as a source data file linked to this manuscript.

### Datasets

Chromatin data (ChIP-seq) from CD4^+^ T cells was downloaded from ENCODE: H3K4me1 (ENCFF112QDR, ENCFF499NFE, ENCFF989BNS), H3K9me3 (ENCFF044NLN, ENCFF736KRZ, ENCFF844IWD, ENCFF929BPC), H3K27ac (ENCFF618IUD, ENCFF862SKP), H3K27me3 (ENCFF124QDD, ENCFF298JKA, ENCFF717ODY), H3K36me3 (ENCFF006VTQ, ENCFF169QYM, ENCFF284PKI, ENCFF5040UW), DNAse (GSM665812, GSM665839, GSM701489, GSM701491). Data was analyzed using Seqmonk (v0.33, http://www.bioinformatics.bbsrc.ac.uk/projects/seqmonk/).

RNA-seq data from CD4^+^ T cells (GSM669617) were used for Figure 6B. We calculated the expression (normalized reads from GSM669617) over all integration sites. Thresholds for expression values (upper 1 /8th, upper quarter, half, and above 0) were set to distinguish five different categories, set as the upper 1 /8th of expression values (high), upper quarter–1/8th (medium), upper half-quarter (low), lower half but above 0 (trace), 0 (silent).

CD4^+^ T cells activation data in Figure 5A was downloaded from GEO (GSE60235).

### Statistical analysis

Significance was analyzed by either paired t-test (GraphPad Prism) or proportion test (standard test for the difference between proportions), also known as a two-proportion z test (https://www.medcalc.org/calc/comparison_of_proportions.php), and specified in the manuscript.

## SUPPLEMENTAL INFORMATION

Supplemental information includes one figure.

## AUTHORS CONTRIBUTION

E.B, M.D, M.AM, L.C, E.V designed the experiments. E.B, M.D, M.AM, L.C, A.G performed the experiments. E.B, M.D, M.AM, I.TS, P.S analyzed the data. E.B, M.D, M.AM, I.TS, P.S, L.C, A.G, S.P, E.V drafted and revised the manuscript.

## ACKNOWLEDGMENTS

We thank Giovanni Maki, Teresa Roberts and John Carroll for graphic preparation, Gary Howard for editorial assistance, and Veronica Fonseca for administrative assistance. E.B. was supported by a post-doctoral fellowship from UCSF CFAR and a CHRP fellowship. M.D was supported by a CIHR 201311MFE-321128-179658. E.V. was supported by funds from NIH 1R01DA030216, 1DP1DA031126, NIH/NIAID R01Ai117864 NIH/NIDA/1R01DA041742-01, NIH/NIDCR/1R01DE026010-01, and 5-31532. We would also like to thank the Gladstone and the Buck Flow Cores. The Gladstone Flow Core was funded by NIH Grants P30AI027763 and S10 RR028962 and by the University of California, San Francisco-Gladstone Institute of Virology and Immunology Center for AIDS Research (CFAR). The standards for PCR methods were made available with help from the University of California San Francisco-Gladstone Institute of Virology & Immunology Center for AIDS Research (CFAR), an NIH-funded program (P30 AI027763), and NIH/NIAIDR21AI129636 for MAM. We thank the amfAR Institute for HIV Cure Research. J.P.S. was supported by the Swedish Research Council (VR2015-02312) and Cancerfonden (CAN2016/576).

## SUPPLEMENTAL FIGURE LEGENDS

**Figure S1.**
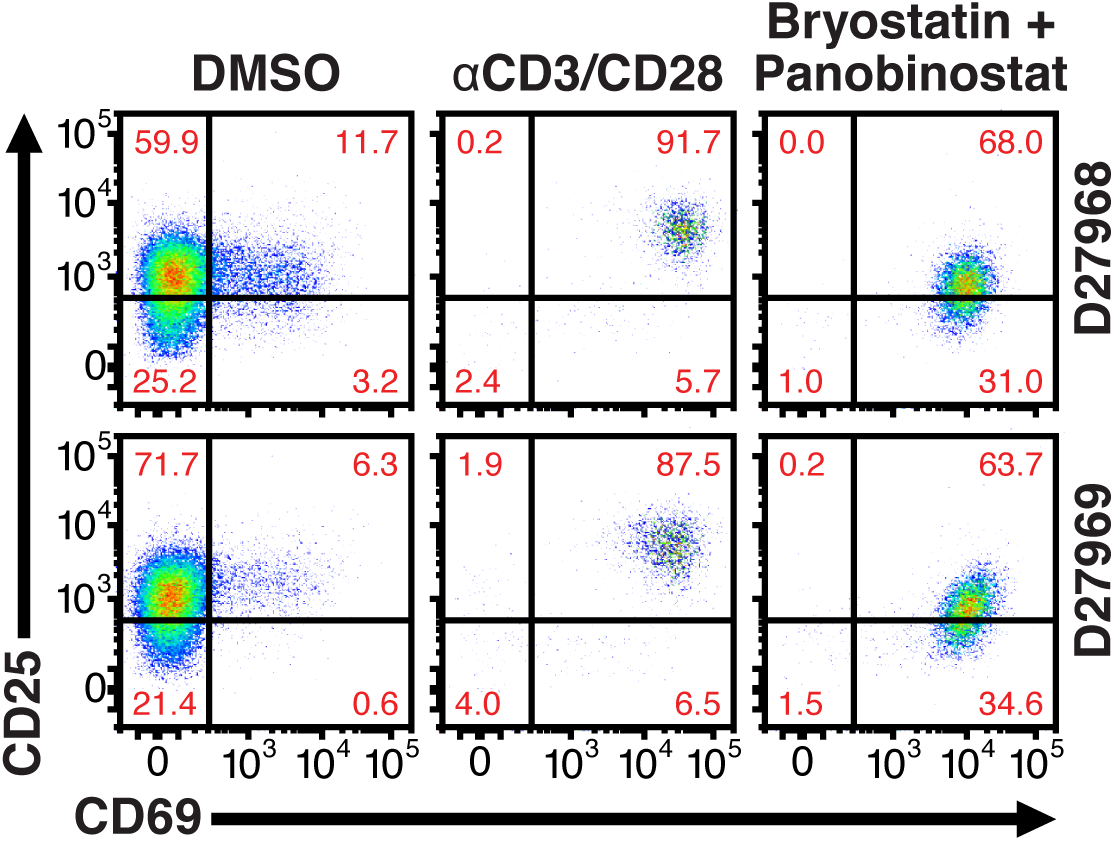
24h treatment effectively activate primary CD4^+^ T cells. Quantification of T-cell activation-associated surface markers after 24hr stimulation. Briefly, CD4^+^ T-cells were purified from blood of two healthy donors and activated for 72h with αCD3/CD28 beads and 100 U/ml IL-2 before infection with HIV_GKO_. Five-days post-infection, latently infected cells (csGFP-mKO2+) were sorted, cultured overnight and stimulated with either αCD3/CD28 or bryostatin + panobinostat in presence of raltegravir. 24h post-treatment, cells were stained for CD25 and CD69 activation markers before performing FACS analysis.

